# Protein disulphide isomerase (PDI) is protective against several types of DNA damage, including that induced by amyotrophic lateral sclerosis-associated mutant TDP-43 in neuronal cells/ in vitro models

**DOI:** 10.1101/2021.08.31.458441

**Authors:** Sina Shadfar, Marta Vidal, Sonam Parakh, Angela S. Laird, Julie D. Atkin

## Abstract

Protein disulphide isomerase (PDI) is a chaperone that catalyses the formation of thiol-disulphide bonds during protein folding. Whilst up-regulation of PDI is a protective mechanism to regulate protein folding, an increasingly wide range of cellular functions have been ascribed to PDI. Originally identified in the endoplasmic reticulum (ER), PDI has now been detected in many cellular locations, including the nucleus. However, its role in this cellular compartment remains undefined. PDI is implicated in multiple diseases, including amyotrophic lateral sclerosis (ALS), a fatal and rapidly progressing neurodegenerative condition affecting motor neurons. Loss of essential proteins from the nucleus is an important feature of ALS. This includes TAR DNA-binding protein-43 (TDP-43), a DNA/RNA binding protein present in a pathological form in the cytoplasm in almost all (97%) ALS cases, that is also mutated in a proportion of familial cases. PDI is protective against disease-relevant phenotypes associated with dysregulation of protein homeostasis (proteostasis) in ALS. DNA damage is also increasingly linked to ALS, which is induced by pathological forms of TDP-43 by impairment of its normal function in the non-homologous end-joining (NHEJ) mechanism of DNA repair. However, it remains unclear whether PDI is protective against DNA damage in ALS. In this study we demonstrate that PDI was protective against several types of DNA damage, induced by either etoposide, hydrogen peroxide (H2O2), or ALS-associated mutant TDP-43^M337V^ in neuronal cells. This was demonstrated using widely used DNA damage markers, phosphorylated H2AX and 53BP1, which is specific for NHEJ. Moreover, we also show that PDI translocates into the nucleus following DNA damage. Here PDI is recruited directly to sites of DNA damage, implying that it has a direct role in DNA repair. This study therefore identifies a novel role of PDI in the nucleus in preventing DNA damage.

## Introduction

Protein disulphide isomerase (PDI) is a multifunctional and highly abundant chaperone that is crucial for protein folding, and the prototype of an extended family of related proteins. It catalyses the formation of disulphide bonds via oxidoreductase activity [1] and it also refolds unfolded or misfolded proteins by chaperone activity [1]. Disulphide bonds play a pivotal role in maintaining the structure of proteins to ensure their performance in many cellular functions. Up-regulation of PDI is a cellular protective mechanism during the unfolded protein response (UPR) following endoplasmic reticulum (ER) stress, when misfolded/unfolded proteins accumulate in the ER [2]. Protein misfolding can lead to the formation of pathological protein aggregates that are associated with many diseases, including neurodegenerative conditions. PDI expression prolongs the survival of mammalian cells, and upregulation of PDI can modulate apoptosis, highlighting its importance in normal cellular function [2–5].

Originally identified in the ER, PDI has now been detected in multiple cellular locations, where an array of cellular functions has been ascribed [6–9]. A growing body of evidence suggests that PDI is found at the surface of several eukaryotic cells [8], where it is implicated in cell-to-cell contact [10], formation of the thrombus on the surface of platelets [11], and the entry of pathogens [12, 13]. PDI has also been detected in the cytoplasm [14, 15], where it redistributes away from its normal ER location via a “protein reflux” system during ER stress [16, 17]. Members of the reticulon family of proteins maintain ER function and can re-distribute PDI away from the ER when overexpressed [18]. In addition, they are implicated in modulating disease progression in mouse models of ALS by a PDI-dependent mechanism [19]. PDI has also been detected in the nucleus, where it anchors DNA loops to the nuclear matrix [9]. In addition, endoplasmic reticulum protein 57 (ERp57), a family member and closest homologue to PDI, has also has been detected in the nucleus, where it was linked to the nuclear import of STAT3 [9]. Nevertheless, the functions of PDI proteins in the nucleus remain poorly defined.

The DNA damage response (DDR) is an important process in the nucleus because cells are under constant attack from both exogenous and endogenous sources, including oxidative stress [20]. The DDR detects and repairs DNA damage to combat these threats. However, unrepaired DNA damage leads to apoptosis or senescence to prevent replicating the damaged genome. Hence the DDR is essential for cellular health and viability, which is particularly important in post-mitotic cells, such as neurons. The most deleterious type of DNA damage is the formation of double-stranded DNA breaks (DSBs), which are primarily repaired by the non-homologous end-joining (NHEJ) DNA repair mechanism in neurons.

Amyotrophic lateral sclerosis (ALS) is a fatal rapidly progressing neurodegenerative disorder affecting motor neurons in the brain, brainstem, and spinal cord, leading to muscle wasting due to denervation. The clinical expression of ALS usually appears in mid-life (between 50-60 years of age), implying that neurons die through cumulative damage to normal cellular mechanisms. The major pathological hallmark of ALS is the formation of misfolded protein inclusions and dysfunction to proteostasis mechanisms are now well described in ALS [21]. PDI has been previously shown to be protective against pathological processes associated with proteostasis in ALS, including protein misfolding, inclusion formation, ER stress, protein trafficking disruption, UPS dysfunction and apoptosis in neuronal cells [14, 22].

Defects in DNA repair, leading to DNA damage, are also increasingly implicated in ALS, and a growing number of proteins with normal functions in DNA repair are now associated with ALS. This includes TAR DNA-binding protein-43 (TDP-43), a DNA/RNA binding protein normally found in the nucleus, that is present in a pathological form in the cytoplasm in almost all (97%) ALS cases. Mutations in TDP-43 are also present in 4-5% of familial forms of ALS cases [23, 24]. Hence TDP-43 is central to neurodegeneration in ALS [25–28]. In the nucleus, TDP-43 normally performs important functions in RNA metabolism and in NHEJ DSB repair [29, 30]. However, pathological forms of TDP-43 lose these normal functions, and mislocalise to the cytoplasm, where they gain aberrant, toxic functions [30]. ALS-associated mutations in TDP-43, including M337V, induce DNA damage and TDP-43 pathology [27, 31]. DNA damage has also been detected in the CNS of patients with sporadic ALS [32–34] and in disease models based on other misfolded proteins in ALS [32, 35–41]. Oxidative stress is another important pathogenic mechanism that can trigger DNA damage, which is also widely implicated in ALS [42, 43].

Given that PDI has been detected in the nucleus [14], we hypothesised that it may also be protective against DNA damage. In this study we demonstrate that PDI is protective against several types of DNA damage, induced by either etoposide, H_2_O_2_, or ALS-associated mutant TDP-43^M337V^ in neuronal cells. Moreover, we also show that PDI translocates into nucleus where it co-localises with DNA damage foci, implying that it has a direct role at DNA damage sites and thus in DNA repair. This study therefore identifies a novel nuclear function for PDI, in preventing DNA damage.

## Materials and Methods

### Constructs

A previously generated pcDNA3.1(+) construct encoding PDI (tagged with V5) was generously provided by Professor Neil Bulleid, University of Glasgow, UK [44]. TDP-43 WT and mutant TDP-43^M337V^, cloned in the pCMV6-AC-GFP vector to allow expression of human TDP-43 with a C-terminal GFP tag, were as previously described [45].

### Cell culture maintenance

Two cell lines were used in this study, Neuro-2a (mouse neuroblastoma) cells, purchased from Cellbank Australia (CODE: 89121404), and NSC-34 (motor neuron-like) cells, which were generously provided by Professor Neil Cashman (University of Toronto, Canada). Cells were cultured in Dulbecco’s Modified Eagle’s Medium high in glucose (DMEM)(Gibco), containing 10% (v/v) heat-inactivated FCS at 37°C, in a 5%, CO_2_ humidified atmosphere. To passage the cells, they were washed with phosphate buffered saline (PBS, 3.2 mM Na_2_HPO_4_, 0.5 mM KH_2_PO_4_, 1.3 mM KCl, 135 mM NaCl, pH 7.4), detached by treatment with 1 mL of Trypsin-EDTA (Gibco), incubated at 37 °C for approximately 2 min then washed with 5 mL of DMEM with 10% FCS to resuspend. Following centrifugation at 1000 g for 5 min, the cell pellet was resuspended in 1 mL of DMEM with 10% FCS, diluted 1:10 in DMEM and counted using a Scepter™ 2.0 Cell Counter (Merk Millipore).

### Immunocytochemistry

Following induction of DNA damage for 30 minutes, cells were washed with 1 x Phosphate-Buffered Saline (PBS) (Invitrogen), and then permeabilised in 0.1% Triton X-100 in PBS for 10 minutes, followed by blocking in 1% BSA in PBS for 1 hour. Cells were then incubated overnight at 4°C with the following primary antibodies at a concentration of 1:500: mouse anti-V5 (1:500, Invitrogen), rabbit anti-γ-H2AX (1:500, Novus Biologicals, NB-100-384), or rabbit anti-53BP1 (1:500, Novus Biologicals, NB100-304). The following secondary antibodies were used: antirabbit Alexa Fluor 488 (1:500, Life Technologies, A11008), anti-mouse Alexa Fluor 594 (1:500, Life Technologies, A21203) or anti-rabbit Alexa Fluor 647 (1:500 Life Technologies, A21245). The nuclei were counterstained with Hoechst 33342 (1:3000, Sigma-Aldrich) and cells were then mounted and imaged. Using fluorescent microscopy, the presence of γH2AX foci was then quantified (as number of foci per 100 cells). Cells containing large, overlapping foci or foci that could not be counted (identified as large foci attached to each other, where it was difficult to determine how many foci were present) were not considered.

### Confocal Microscopy and Image acquisition

Cells were photographed with 63x/na=1.4 or 100x/na=1.46 objectives on a Zeiss LSM 880 inverted confocal laser-scanning microscope, equipped with a LSM-TPMT camera (Zeiss). In multichannel imaging, photomultiplier sensitivities and offsets were set to a level at which bleed through effects from one channel to another were negligible.

### Western Blotting

Soluble protein cellular lysates (20μg) were analysed by 4-15% gradient SDS-PAGE and blotted onto nitrocellulose membranes (Bio-Rad). Membranes were blocked with 5% skim milk in 1 x PBS (Invitrogen) for 1.5 hours and incubated with the respective primary antibodies diluted in blocking buffer for 24 hours at 4°C; anti-γ-H2AX (1:1000, Novus Biologicals, NB-100-384), anti-P4HB (PDI) (1:2000, abcam, ab3672) and GAPDH (1:4000, Proteintech 60004-Ig).

Twenty-four-hours post-incubation, unbound primary antibodies were removed by washing in TBST and the blots were incubated with HRP-conjugated secondary antibodies for 1 hour at room temperature: goat anti-mouse IgG and IgM HRP conjugated antibodies (1:4000, Merck EMD Millipore, Ap130P), or goat anti-rabbit IgG antibody, peroxidase-conjugated (1:4000, Merck EMD Millipore, Ap132P). Immunoreactivity was revealed using the Clarity™ ECL Western Blotting Substrate kit (BioRad) and images were obtained using a BioRad ChemiDoc MP system, using Image Lab™ software (BioRad). The intensity of each band relative to GAPDH was quantified using ImageJ software (v. 1.47; National Institutes of Health).

### Subcellular fractionation

Neuro-2a cells were transfected with either the PDI construct or pcDNA3.1(+) empty vector (EV). Twenty-four hours post-transfection, cells were treated with either 13.5μM etoposide or DMSO for 30 minutes and washed with ice-cold PBS. Then, cells were lysed with fractionation buffer (20mM HEPES (pH 7.4), 10mM KCI, 1.5 mM MgCl2, 1 mM EDTA). Following centrifugation (720 g for 5 min), the supernatant, containing the cytoplasm, membrane, and mitochondrial fractions, was transferred into fresh tubes. The pellet fraction, containing the nuclei, was washed with 500 μl fractionation buffer, and centrifuged again twice, at 720 *g* for 10 minutes. Afterwards, the pellet was resuspended in 200 μl RIPA buffer, with 0.1% SDS followed by brief sonication (10 second on ice), to shear genomic DNA and homogenize the lysate. The primary supernatant was centrifuged at 10,000 g (8,000 rpm) for 5 min. The resulting supernatant containing the cytoplasm was concentrated by centrifugation and frozen at −20°C until required.

The total amount of protein in each sample was quantified using a Pierce BCA Protein Assay Kit (Thermoscientific), following the manufacturer’s instructions. Concentrated protein samples (10-20μg) were separated on 7.5% or 4-15% (BioRad) SDS-PAGE gels. Proteins were then transferred onto nitrocellulose membranes according to the manufacturer’s instructions (BioRad) and blotted as above using polyclonal mouse anti-P4HB (PDI) (1:2000, abcam, ab3672), antirabbit Lamin B (1:3000, Abcam, ab16048) or GAPDH (1:4000, Proteintech, 60004-Ig) antibodies. After rinsing, the blots were incubated in peroxidase-conjugated secondary antibodies (1:2000; Millipore) for one hour at room temperature.

### Statistics

Data are presented as mean value ± standard error of the mean (SEM). Statistical comparisons between group means were performed and graphs were generated using GraphPad Prism 7.02 software (Graph Pad software, Inc.). A t-test or a one-way ANOVA, followed by a *post hoc* Tukey test for multiple comparisons, was used when justified. The significance threshold was set at *p* = 0.05. The number of independent repeats (n), the statistical test used for comparison and the statistical significance (*p* values) are specified for each figure panel in the representative figure legend.

## Results

### PDI inhibits the formation of DNA damage foci induced by etoposide

To investigate whether PDI is protective against DNA damage *in vitro*, Neuro-2a cells, a mouse neuroblastoma cell line, were transfected with PDI tagged with V5 or pcDNA3.1(+) empty vector. At 24 hours post-transfection, the cells were treated with 13.5 μM etoposide to induce DNA damage, or DMSO as a control. Etoposide induces single and double stranded breaks in DNA by targeting DNA topoisomerase II [46, 47]. Phosphorylation of H2AX at Ser 139 (γ-H2AX) and the formation of discrete γ-H2AX foci is a widely used and sensitive DNA damage marker [48]. Hence, immunocytochemistry was then performed using both anti-γH2AX (green) and anti-V5 (red) antibodies. Quantification using fluorescence microscopy revealed that DNA damage, quantified by the number of γH2AX foci produced, increased significantly in all etoposide treated groups compared to DMSO controls, confirming the induction of DNA damage. Moreover, significantly fewer γH2AX foci were formed in etoposide-treated cells expressing PDI, compared to both untransfected (UT) and EV transfected cells **(Figure 1, A, B ****p < 0.0001)**. Hence this implies that PDI is protective against induction of DNA damage induced by etoposide.

**Figure 1.**
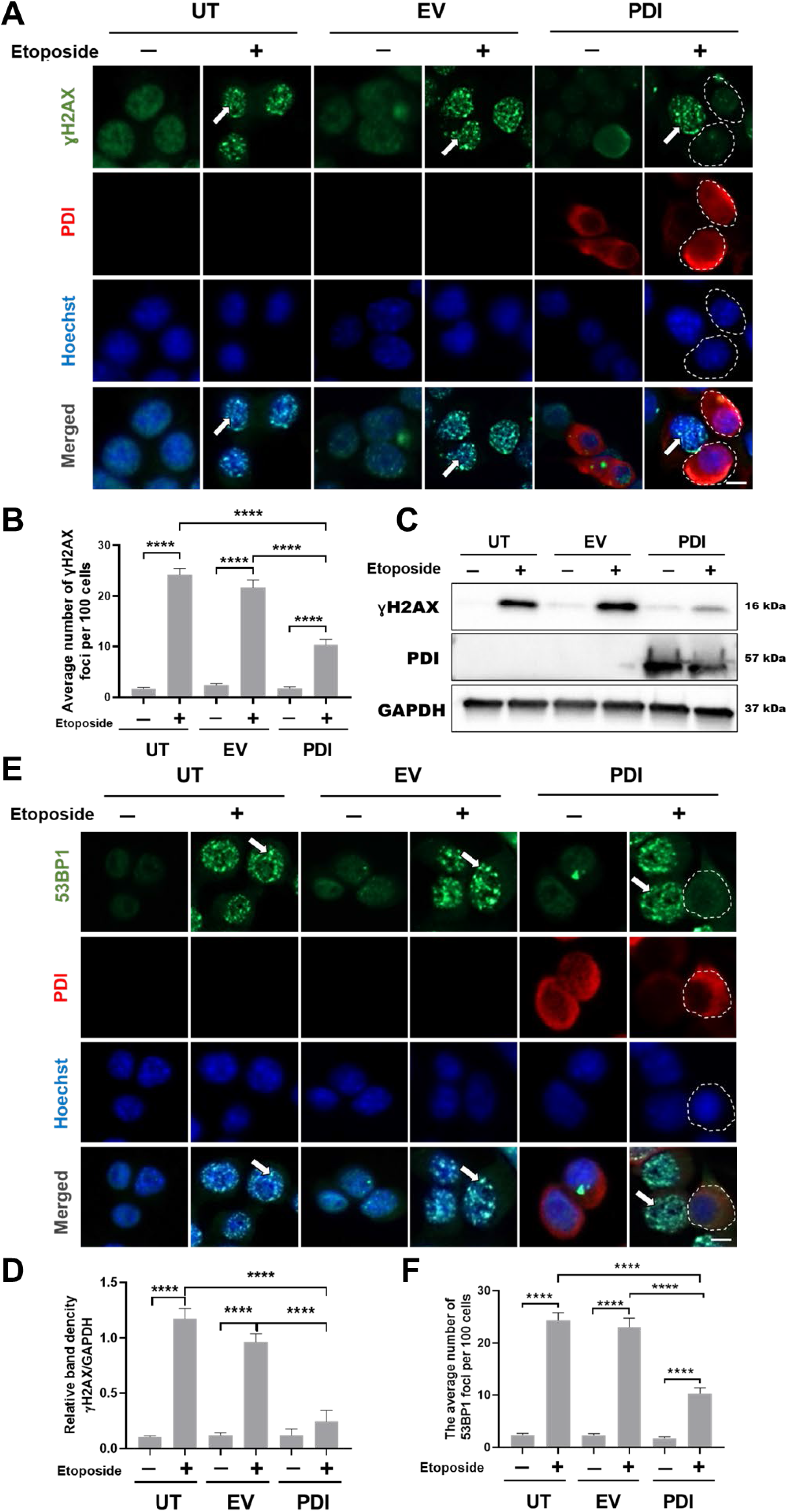
PDI is protective against DNA damage induced by etoposide in Neuro-2a cells. **(A, E)** Neuro-2acells were transfected for 24 hours with constructs encoding either pcDNA empty vector (EV) or PDI tagged with V5. After 30 minutes treatment with either etoposide or DMSO only, immunocytochemistry was performed using an anti-V5 antibody (red) and antibodies against either **γ**H2AX (A) or 53BP1 (E) (both green). Nuclei were stained with Hoechst stain (blue). Scale bar: 10μm. Arrows represent DNA damage foci, **γ**H2AX (A) or 53BP1 (E). **(B, F)** Quantification of the average number of foci per 100 cells shown in (A) or (E) (n=3). Over-expression of PDI significantly reduced the number of DNA damage foci, γH2AX or 53BP1 in Neuro-2a cells following etoposide induced DNA damage**. (C)** Western blotting for **γ**H2AX, PDI and GAPDH (as a loading control) in Neuro-2a cells transfected with pcDNA empty vector or PDI tagged with V5, following 30 minutes etoposide treatment. Blot confirms the overexpression of PDI. **(D)** Quantification of **γ**H2AX levels in the blots in (C). Expression of **γ**H2AX was significantly reduced in PDI overexpressing cells following etoposide treatment, compared to untransfected (UT) or empty vector-cells (EV). The graph depicts the band intensity of **γ**H2AX relative to GAPDH. **** p < 0.0001, One-way ANOVA followed by Tukey’s multiple comparison post-hoc test. All values represent mean ± SEM.

Next, western blotting was performed to examine the levels of γH2AX (which becomes up-regulated during DNA damage) following PDI expression in etoposide treated cells. Quantification using densitometry revealed induction of DNA damage following etoposide treatment as expected, compared to DMSO. However, lower levels of γH2AX were present in cells expressing PDI following etoposide treatment, compared to the EV and UT cells (four-fold) **(Figure 1 C, D, ****p < 0.0001)**. This data therefore provides further evidence that expression of PDI protects against DNA damage induced by etoposide in Neuro-2a cells.

Secondly, another marker of the DDR, p53-binding protein 1 (53BP1), was examined. 53BP1 is a regulator of DSB repair and it is recruited to the nucleus following DNA damage, similar to γH2AX [49], where it forms specific foci at sites of DNA damage. Fluorescence microscopy and quantification revealed that the number of 53BP1 DNA damage foci increased significantly after 30 minutes of etoposide treatment in all groups, compared to the respective DMSO controls, confirming induction of DNA damage. Importantly, there were significantly fewer 53BP1 foci in etoposide treated cells expressing PDI compared to UT and EV controls **(Figure 1E, F, ****p <0.0001)**. Thus, these findings further confirm a protective role for PDI against etoposide-induced DNA damage. In addition, because 53BP1 is specific for the NHEJ mechanism of DNA repair, this implies that PDI is protective against the formation of DSBs during NHEJ repair.

Similar experiments were performed in another cell line to confirm that these results were not specific to Neuro-2a cells only. The NSC-34 cell line is a hybrid of spinal cord motor neurons and a mouse neuroblastoma cell line, N18TG2 [50]. NSC-34 cells display several physiological and morphological properties of motor neurons, and therefore they are also relevant to ALS [50]. Hence, NSC-34 cells were transfected with either PDI tagged with V5 or pcDNA3.1(+) EV. At 24-hour post-transfection, cells were treated for 30 minutes with either 13.5 μM of etoposide or DMSO only in the control group. Immunocytochemistry using anti-γH2AX (green) and anti-V5 antibodies (red) was performed. Fluorescence microscopy and quantification confirmed induction of DNA damage in all etoposide-treated groups compared to DMSO controls **(Supplementary Figure 1A, B, ****p <0.0001)**. Moreover, significantly fewer γH2AX DNA damage foci were formed after etoposide treatment in PDI overexpressing cells, compared to the untransfected and EV-transfected cells **(Supplementary Figure 1A ****p <0.0001)**. These results confirm that PDI is protective against DNA damage in NSC-34 cells.

Similarly, the number of 53BP1 DNA damage foci was next examined in NSC-34 cells overexpressing PDI compared to controls. Induction of DNA damage was confirmed by the presence of significantly more 53BP1 foci in etoposide-treated cells compared to DMSO controls **(Supplementary Figure 1 C, D)**. Furthermore, in comparison to the untransfected and EV transfected cells, PDI overexpressing cells displayed significantly fewer DNA damage foci following etoposide treatment **(Supplementary Figure 1D, ****p ≤0.0001)**. Thus, these results confirm that PDI expression inhibits DNA damage in NSC-34 cells, following etoposide treatment, thus validating the results in Neuro2a cells.

### PDI inhibits DNA damage foci induced by H_2_O_2_ in Neuro-2a cells

Since PDI was protective against DNA damage induced by etoposide, we next investigated whether PDI was also protective against other forms of DNA damage. Reactive oxygen species (ROS) such as H_2_O_2_ are the main source of oxidative stress in living organisms. They are highly reactive towards DNA [51], can inhibit DNA repair enzymes [52, 53] and are widely implicated in several neurodegenerative disorders, including ALS [51]. Hence, we next used H_2_O_2_ to induce oxidative DNA damage, given its relevance to cellular function and ALS [54].

Neuro-2a cells were transfected with either PDI tagged with V5 or pcDNA3.1(+) EV. At 24hour post-transfection, cells were treated with either 100 μM H_2_O_2_ or DMSO for 1 hour. Immunocytochemistry and fluorescence microscopy was then performed using anti-V5 and anti-γH2AX antibodies. A significant increase in the number of γH2AX DNA damage foci following H_2_O_2_ treatment in all etoposide groups relative to DMSO only controls, confirmed induction of DNA damage **(Figure 2 A, B, ****p <0.0001)**. In contrast, following H_2_O_2_ treatment, significantly fewer γH2AX DNA foci were formed in PDI overexpressing cells, compared to untransfected and EV-transfected cells **(Figure 2B, p <0.0001)**. Hence, these results show that PDI is also protective against H_2_O_2_-induced DNA damage in Neuro-2a cells.

**Figure 2.**
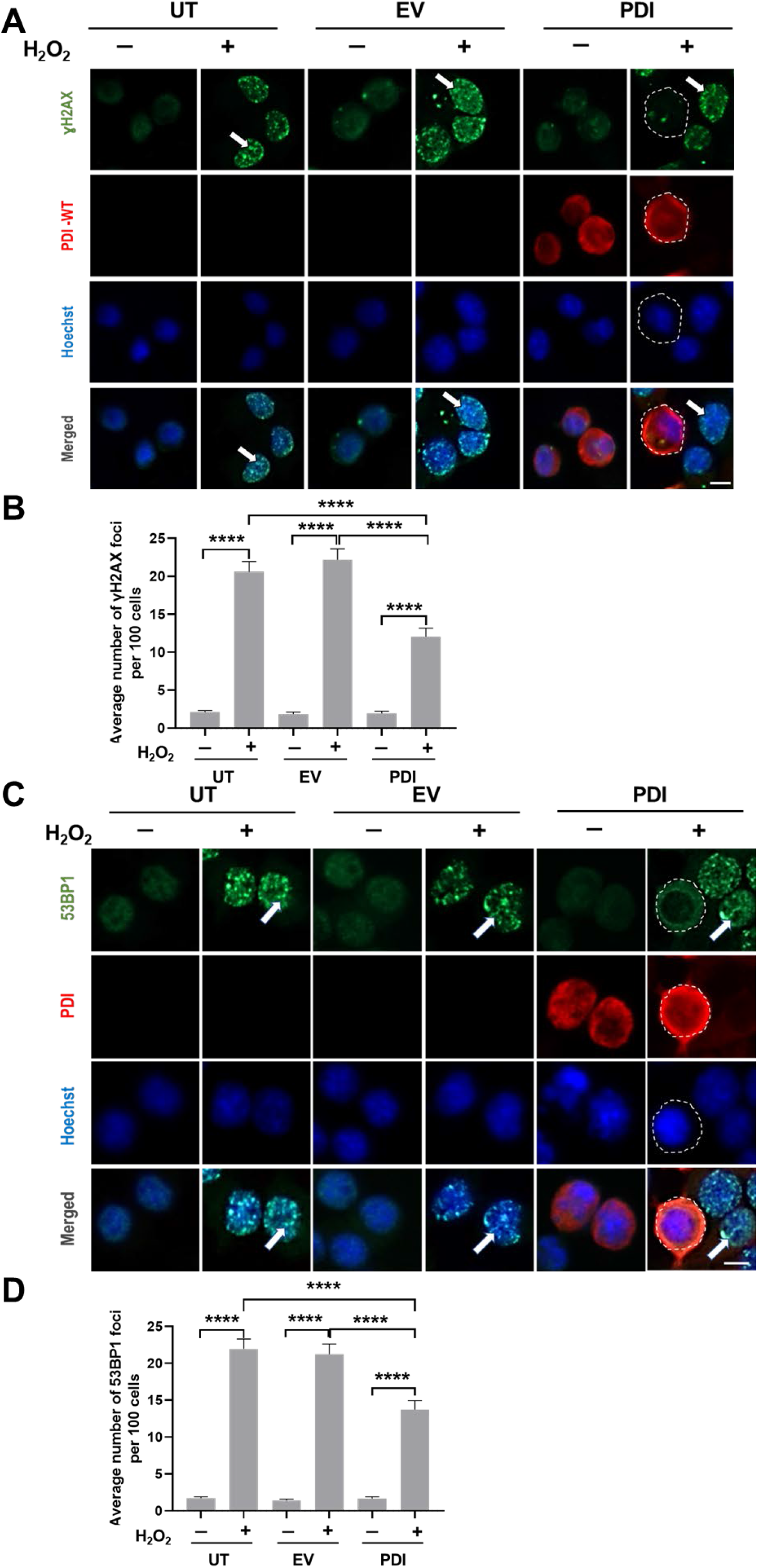
PDI inhibits the formation of DNA damage foci induced by H_2_O_2_ in neuronal cells Neuro-2a cells. **(A, C)** Neuro-2a cells overexpressing PDI tagged with V5 (red), or empty vector were subjected to immunocytochemistry using antibodies against **γ**H2AX (A) or 53BP1 (C) (both green), and V5 (red), following 1-hour H_2_O_2_ treatment (100μM). Nuclei were stained with Hoechst (blue). Arrows represent 53BP1 foci; Scale bar: 10 μm. **(B, D)** Quantification of the average number of DNA damage foci per 100 cells shown in (A). PDI over-expression significantly reduced the number of **γ**H2AX and 53BP1 foci induced by H_2_O_2_ in Neuro-2a cells. n=3 **** p <0.0001, one-way ANOVA followed by Tukey’s multiple comparison post-hoc test. All values represent mean ± SEM

To further confirm these results, the formation of 53BP1 foci was examined next. Immunocytochemical analysis and fluorescence microscopy demonstrated that significantly more 53BP1 DNA damage foci were present in cells treated with H_2_O_2_ for all groups, compared to DMSO controls **(Figure 2A, B, ****p ≤0.0001)**. Importantly, PDI expressing cells displayed significantly fewer 53BP1 DNA damage foci than untransfected and empty pcDNA3.1(+) vector-transfected cells treated with H_2_O_2_ **(Figure 2B ****p ≤0.0001)**. Thus, these results confirm that PDI expression inhibits the formation of 53BP1 DNA damage foci following etoposide treatment, providing additional evidence that PDI is protective against the formation of DSBs during NHEJ repair.

### PDI overexpression is protective against mutant TDP-43^M337V^ induced DNA damage

We next investigated whether PDI was protective against DNA damage induced by mutant TDP-43^M337V^. We previously established that TDP-43^Q331K^ and TDP-43^A315T^ induce DNA damage in NSC-34 cells [27]. Similarly, we also demonstrated that fibroblasts obtained from mutant TDP-43^M337V^ ALS patients display more DNA damage than control fibroblasts, although this was not shown in cell lines [27]. Hence, we first confirmed that TDP-43^M337V^ induces DNA damage in Neuro2a cells transfected as above.

Following immunocytochemistry for γH2AX as above, followed by fluorescence microscopy and quantification, γH2AX foci were increased significantly following etoposide, compared to DMSO, treatment, confirming induction of DNA damage **(Figure 3 A, B, ****p <0.0001)**. As in previous studies, TDP-43 WT overexpressing cells displayed significantly fewer DNA damage foci, compared to control-cells transfected with EV **(Figure 3 A, B, ****p <0.0001)**, consistent with its recently ascribed DNA repair function [27, 30]. In contrast, there was a significant increase in the number of γH2AX DNA damage foci in TDP-43^M337V^ expressing cells, compared to TDP-43 WT expressing cells **(Figure 3 A, B, ****p <0.0001).** Furthermore, there was no significant difference in the number of foci in TDP-43^M337V^ expressing cells, compared to EV transfected cells **(Figure 3 A, B, *p >0.05)**. This finding confirms that ALS-associated mutant TDP-43^M337V^ lacks the normal function of TDP-43 WT in DNA repair, leading to the accumulation of DNA damage, consistent with the findings obtained previously using M337V fibroblasts.

**Figure 3.**
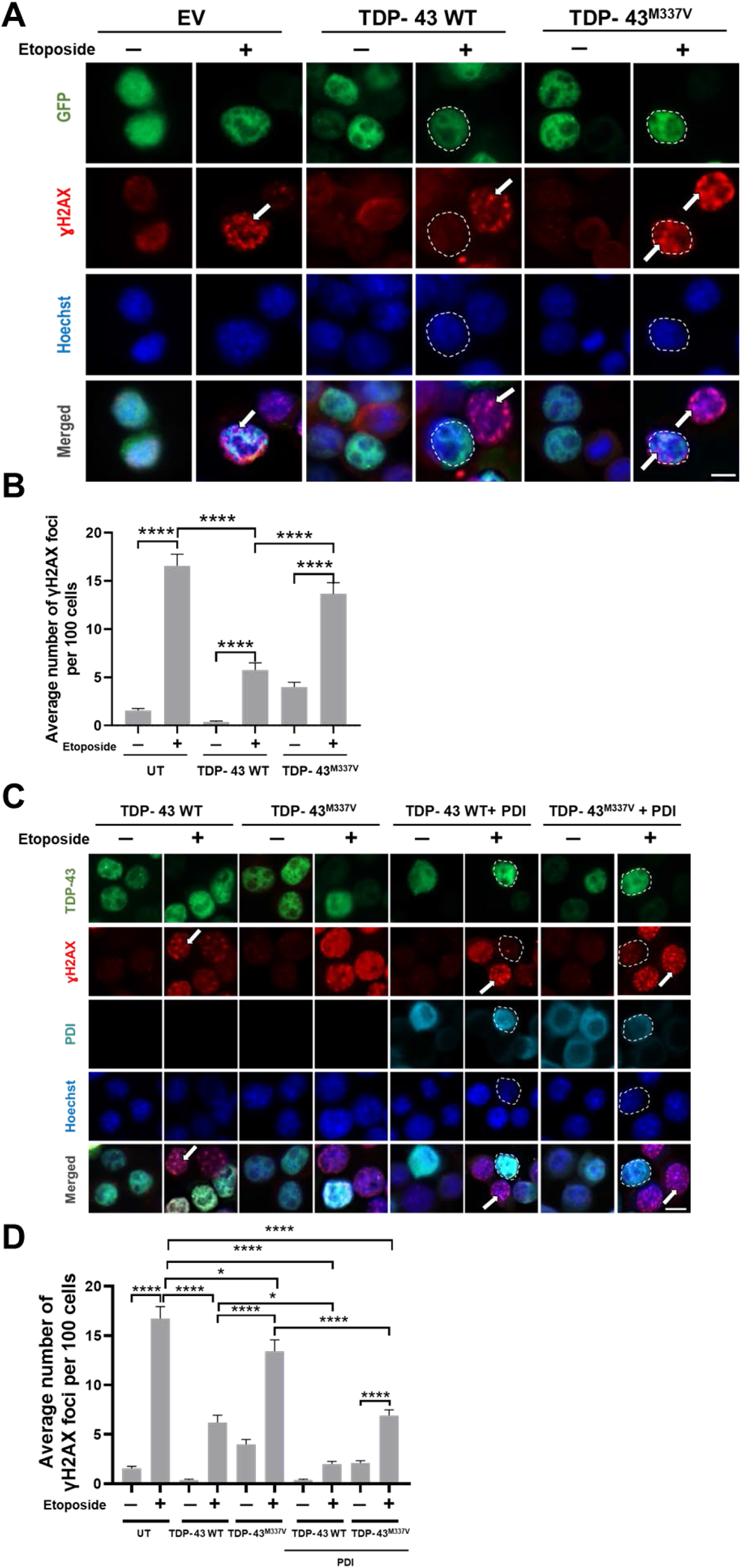
PDI-WT reduces the number of DNA damage foci induced by etoposide in mutant TDP43^M337V^ in Neuro2a cells. A) Neuro-2a cells were transfected with either GFP-tagged (green) TDP-43 WT, TDP-43^M337V^ or empty vector EGFP only. Cells were treated with 13.5μM etoposide for 30 minutes, at 24 hours post-transfection. Immunocytochemistry was performed using an anti-γH2AX antibody (red) and the nuclei were stained with Hoechst (blue). Arrows represent γH2AX foci; Scale bar: 10 μm. B) Quantification of the average number of foci per 100 transfected cells shown in (A). The number of etoposide-induced γH2AX DNA damage foci decreased significantly upon TDP-43 WT expression compared to EV and UT cells, but not for mutant TDP-43^M337V^, indicating that TDP-43^M337V^ lacks the normal protective function of TDP-43 WT C) Neuro-2a cells were co-transfected with either EGFP-tagged TDP-43 WT, TDP43^M337V^ (green) or empty EGFP expressing vector, and PDI tagged with V5 or empty V5 vector. Cells were treated with 13.5μM etoposide or DMSO for 30 minutes at 24 hours post-transfection. Immunocytochemistry was then performed using anti-**γ**H2AX (red) and anti-V5 antibodies (turquoise). The nuclei were stained with Hoechst (blue). Scale bar: 10 μm. D) Quantification of the average number of **γ**H2AX foci per 100 WT or mutant TDP-43^M337V^ (or EGFP for EV) expressing cells in (A). n=3, * p < 0.05, **** p <0.0001; Only cells co-expressing both PDI and TDP-43 (or EGFP) together in the same cell, were included in the analysis. One-way ANOVA followed by Tukey’s multiple comparison post-hoc test. All values represent mean ± SEM.

Having established that mutant TDP-43^M337V^ lacks the normal protective function of TDP-43 WT, and hence that more DNA damage is induced following etoposide treatment in TDP-43^M337V^ expressing cells, it was next examined whether PDI was protective against mutant TDP-43 ^M337V^ induced DNA damage. Cells were transfected with TDP-43 constructs and PDI-V5 construct or with EV. Immunocytochemistry and fluorescent microscopy revealed that in all groups, significantly more γH2AX foci were present following etoposide treatment for 30 minutes, confirming induction of DNA damage. Again, significantly fewer DNA damage foci were present in TDP-43 WT overexpressing cells compared to EV, confirming its protective activity in DNA repair **(Figure 3 C, D, ****p <0.0001)**. However mutant TDP43^M337V^ cells formed significantly more foci than TDP-43 WT expressing cells, confirming that this mutant induces DNA damage in cell lines, similar to other TDP-43 mutants **(Figure 3, C, D, ****p <0.0001)**. However, significantly fewer γH2AX foci were formed in cells co-expressing TDP43^M337V^ and PDI, compared to those cells expressing TDP43^M337V^ with empty V5 vector **(Figure 3 A, B *p < 0.05)**. These data therefore imply that PDI is protective against DNA damage induced by TDP43^M337V^. They also suggest that PDI is protective by a mechanism distinct from TDP-43 WT, because significantly fewer γH2AX foci were formed in cells expressing TDP-43 WT and PDI compared to TDP-43 WT with EV **(Figure 3 C, D, *p < 0.05).**

### PDI translocates into the nucleus following induction of DNA damage

The results described above reveal that PDI is protective against DNA damage induced by etoposide, H_2_O_2_ and ALS-associated mutant TDP-43. However, it is unclear in which cellular location this protective activity is mediated because DNA damage occurs in the nucleus, whereas PDI is conventionally considered to be localised in the ER. Hence, the subcellular localisation of PDI following etoposide treatment was investigated, to determine if PDI translocates into the nucleus following induction of DNA damage. Endogenous PDI was examined first rather than over-expressed PDI so that the normal physiological response could be examined.

We investigated the localisation of endogenous PDI in untransfected cells by immunohistochemistry using antibodies against PDI and γH2AX, following induction of DNA damage using etoposide. In etoposide treated cells, PDI appeared to form foci in the nucleus, but these structures were rare in DMSO treated cells. Quantification revealed these nuclear PDI foci were significantly more abundant in etoposide treated cells, compared to DMSO only cells (106 foci, 1 foci respectively per 100 cells, **Figure 4A, B, ***p < 0.001)**. This finding suggests that following DNA damage induction, PDI translocates into the nucleus, presumably from its primary location, the ER. This implies that PDI has a direct role in the nucleus in response to DNA damage.

**Figure 4.**
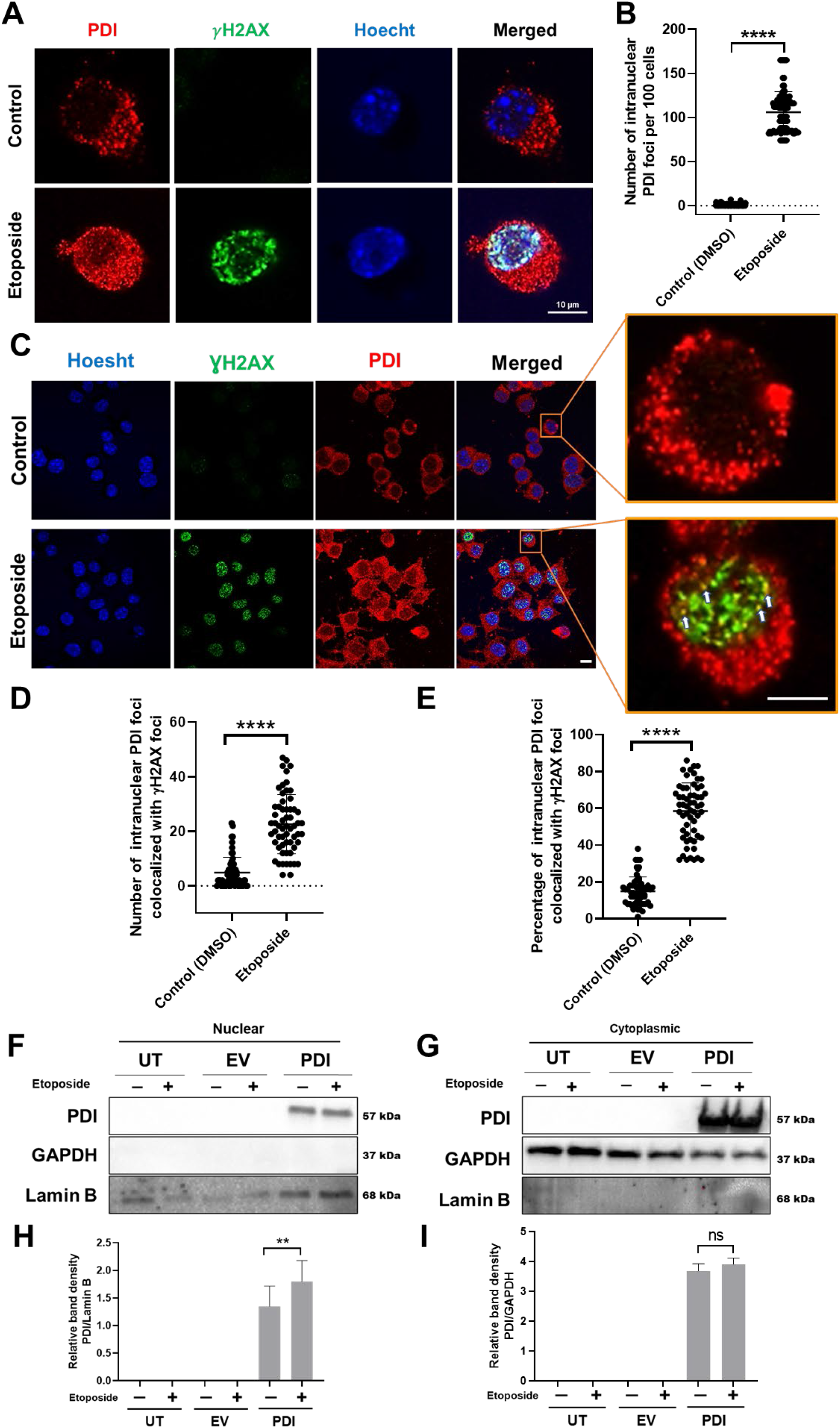
PDI is recruited to DNA damage foci in the nucleus following etoposide treatment. **(A)** Immunocytochemistry to examine endogenous PDI expression following 30 minutes of etoposide or DMSO treatment in Neuro-2a cells, using anti-**γ**H2AX (green) and anti-PDI antibodies (red) using confocal imaging. Nuclei were stained with Hoechst (blue). **(B)** Quantification of the average number of PDI foci within the nucleus per 100 cells in (A). PDI foci were abundant in the nucleus of etoposide treated cells but rare in DMSO treated cells, n=3. Scale bar: 10 μm.**p< 0.01 ****p< 0.0001 (unpaired t test). **(C)** Immunocytochemistry following 30 minutes of etoposide or DMSO treatment in Neuro-2a cells expressing PDI-V5, using anti-**γ**H2AX (green) and anti-PDI antibodies (red) and confocal imaging. Nuclei were stained with Hoechst (blue). Arrows represent colocalization of intranuclear PDI with **γ**H2AX foci. **(D)** Quantification of the number of PDI foci colocalized with **γ**H2AX foci per 100 cells in (C). n=3, ****p < 0.0001. One-way ANOVA followed by Tukey’s multiple comparison post-hoc test. **(E)** Quantification of the percentage of intracelleular PDI foci colocalized with **γ**H2AX foci per 100 cells in (C). n=3, ****p < 0.0001. One-way ANOVA followed by Tukey’s multiple comparison post-hoc test. **(F, G)** Western blotting of nuclear (F) and cytoplasmic (G) fractions prepared from lysates of cells expressing empty vector or V5-tagged PDI, or untransfected cells, using an anti-PDI antibody and antibodies against lamin B and GAPDH as markers of the nucleus and GAPDH respectively. **(H, I)** Quantification of PDI levels in the nuclear and cytoplasmic fractions of the blots shown in (F, G), following etoposide or DMSO treatment. The graph depicts the relative band intensity by densitometry of PDI relative to lamin B in the nuclear fractions, and of PDI to GAPDH in the cytoplasmic fractions. The levels of PDI in the nucleus increased significantly following etoposide treatment. n=3, **p < 0.001, ns = non-significant. One-way ANOVA followed by Tukey’s multiple comparison post-hoc test. All values represent mean ± SEM.

We next examined the localisation of overexpressed PDI so that the protective effects of PDI described above could be related to its presence or absence from the nucleus. As above, Neuro-2a cells were transfected with either pcDNA3.1(+) empty vector or V5-tagged PDI and at 24hours post transfection, they were treated with etoposide (or DMSO) for 30 minutes with untransfected cells. Following immunocytochemistry for γH2AX and V5, the presence of PDI at sites of DNA damage was examined by quantifying the co-localisation of PDI and γH2AX using Image J. These analyses demonstrated that 57.4% of the intra-nuclear PDI foci colocalized with γ-H2AX following etoposide treatment, whereas only 12.73% intranuclear PDI foci colocalized with γ-H2AX in the DMSO-treated group **(Figure 4C, D, E, ****p < 0.0001)**. These results imply that overexpressed PDI is recruited specifically to γH2AX foci and hence sites of DNA damage following etoposide treatment. To further confirm these results, cell lysates were prepared from cells transfected as above, and subcellular fractionation was performed to generate the nuclear and cytoplasmic fractions. Western blotting of the nuclear and cytoplasmic fractions using anti-PDI, anti-lamin-B (as a nuclear marker) and anti-GAPDH (as a cytoplasmic marker) antibodies was performed. The lack of GAPDH reactivity in this fraction confirmed that there was little contamination from the cytoplasm. Moreover, these analyses revealed that there was a significant increase in the levels of PDI in the nuclear fraction following etoposide treatment, compared to DMSO treated cells **(Figure 4 F, H, **p < 0.01)**. In contrast, there was no statistically significant difference in the levels of PDI in the cytoplasm following etoposide treatment **(Figure 4G, I, p > 0.05**). Hence these data confirm that both endogenous and overexpressed PDI become recruited to the nucleus following DNA damage, where they co-localise with γH2AX foci.

## Discussion

In this study we describe a novel protective role for PDI against DNA damage. Over-expression of PDI in neuronal cell lines inhibited DNA damage, induced by either etoposide, H_2_O_2_, or ALS-associated mutant TDP-43^M337V^. Moreover, we also provide the first evidence that PDI translocates into nucleus following DNA damage, where it co-localises with γH2AX foci, implying it has a direct role in DNA repair mechanisms.

Each cell receives numerous DNA injuries every day, from both exogenous and endogenous sources, including oxidative stress, topological changes to DNA, and the presence of misfolded proteins [55]. Protection against genetic integrity is essential for cellular and human health [55]. Neurons are post-mitotic cells with high metabolic rates; hence they are more susceptible to DNA damage compared to other cell types. Furthermore, neurons are highly susceptible to oxidative stress, which is a major source of DNA damage [56]. Here we show that PDI overexpression protects neuronal cells against DNA damage, including that induced by oxidative stress and a mutant protein central to neurodegeneration, using both immunocytochemistry and western blotting.

We used two important and widely markers of the DDR in this study: γH2AX and 53BP1 [49, 57–60]. Phosphorylation of the Ser-139 residue of the core histone protein H2AX is one of the earliest cellular responses to the presence of DNA DSBs [61]. Detection of this phosphorylation event is recognised as a highly specific and sensitive molecular marker for monitoring DNA damage [48, 62]. Hence these results imply that PDI is protective against the most cytotoxic type of DNA lesions, DSBs. One role of γH2AX, specifically during the NHEJ mechanism of DSB repair, is to recruit 53BP1 to the vicinity of DSB sites [63, 64]. 53BP1 is an important regulator of the cellular response to DSBs during NHEJ, and it promotes the ligation of distal

DNA ends. Hence our results imply that PDI is active in the NHEJ mechanism of DNA repair, either directly or indirectly. The mechanism for this activity remains unclear, although it could relate to the oxidoreductase activity of PDI given that the DDR is regulated by redox processes [65, 66]. However further studies are required to examine the specific mechanisms involved.

Another possibility is that PDI is involved in the cross-talk between the DNA damage and ER stress. Expression of PDI is induced during the UPR [67, 68], and whilst not well characterised, a link between DNA damage and ER stress has been previously reported [69]. ER stress sensitizes cells to genomic damage [70] and induces apoptosis through p53 activation [71]. Conversely, DNA damage initiates tubular extension of the ER dependent upon activation of p53 to induce apoptosis by facilitating ER-mitochondrial signalling [69].

In this study, DNA damage was induced by three distinct mechanisms and PDI was protective against all three processes, implying it has broad activity in the DDR. Etoposide is an inhibitor of topoisomerase II, which regulates the topological state of DNA and thus manages tangles and supercoils, resulting in the generation of DSBs. In contrast, H_2_O_2_ treatment induces ROS, triggering oxidative DNA damage. Thirdly, DNA damage was induced by expression of mutant TDP-43^M337V^. We and others previously found that TDP-43 WT overexpression is protective against etoposide-induced DNA damage [27, 72], revealing that TDP-43 is a DNA repair protein that functions in NHEJ [27, 30, 73]. However, the ALS-associated TDP-43 mutants lack this activity, inducing DNA damage. Hence, the protective function of PDI against mutant TDP-43 provides further evidence that its defensive activity in DNA damage involves the NHEJ pathway of DNA repair. Interestingly, however, in the current study we showed that co-expression of PDI with TDP-43 WT is significantly more protective against etoposide challenge than TDP-43 expression alone. This implies that TDP-43 and PDI are protective by distinct, rather than the same, molecular mechanisms. TDP-43 is also known to be recruited specifically to sites of DNA damage [72], where it acts as a scaffold for the recruitment of the XRCC4-DNA ligase 4 complex, the major DNA ligase involved in NHEJ [74]. Hence, it is likely that PDI acts via a different process during NHEJ.

Although PDI is primarily located in the ER, it leaves the ER in some circumstances [8], and its localisation in the cytoplasm and cell surface are now well described [8, 9, 75]. In the current study, PDI was found to translocate to the nucleus, and more PDI was present in this location following etoposide treatment, compared to DMSO only. Here we detected the nuclear presence of both endogenous PDI, as a physiological response, and over-expressed PDI, relating its specific protective activity to localisation in the nucleus. Moreover, PDI was found to localise to discrete DNA damage foci in the nucleus following treatment. Hence these findings reveal that PDI translocates directly to DNA damage sites following etoposide treatment, implying that PDI exerts its protective activity here. The identity of the PDI foci remains unclear however and should be investigated further.

Previously the formation of protein complexes, containing PDI and ERp57 in the nucleus interacting with DNA, have been detected [9]. Also PDI can be cross-linked to DNA in the nucleus of HeLa cells, where it anchors to the DNA-nuclear matrix [7]. PDI also initiates the binding of transcription factors nuclear factor-κB (NF-κB) and AP-1 to DNA [76]. It is well established that together with p53, NF-κB alters transactivation of several genes which participate in the DDR [77]. In addition, there is functional cross-talk between the Nrf2 and NF-κB pathways, which regulate cellular responses to oxidative stress and inflammation, respectively. In addition, a link between PDI and DNA damage in relation to cancer was previously postulated. Accumulating evidence suggests that PDI is involved in the growth, metastasis, and survival of multiple types of cancer cells [78, 79], which was associated with DNA damage [80]. Our findings revealing a novel function of PDI in response to DNA damage in the nucleus are therefore consistent with earlier studies describing PDI family members in this location.

Previously a role for cytoplasmic heat shock protein (Hsp) chaperones Hsp70 and Hsp90 in the DDR to DSBs was demonstrated [81]. Inhibition of Hsp90 with 17-(allylamino)-17-demethoxygeldanamycin (17-AAG), leads to loss of ataxia telangiectasia mutated (ATM) activity, a core DNA repair component. [82]. ATM is activated by DSBs by the homologous recombination (HR) repair pathway, although this is thought to be absent from neurons. Hsp70 and Hsp27 are also associated with the base excision repair (BER) pathway, which removes damaged (oxidized or alkylated) bases generated by ROS, via DNA glycosylase (UDG) and human AP endonuclease (HAP1) in HeLa cells [83]. Hsp 70 and Hsp 90 are normally localised in the cytoplasm but are known to be imported into the nucleus following the induction of oxidative DNA damage using H_2_O_2_ or 8-hydroxyguanosine (8-OH-dG)[84], demonstrating that cellular chaperones can re-locate in response to DNA damage

We previously showed that PDI was protective against several pathological features related to proteostasis induced by mutant TDP-43 in ALS, including inclusion formation, mislocalisation to the cytoplasm, ER stress, trafficking disruption, and apoptosis in neuronal cells [14, 22]. Novel roles for PDI proteins were also recently identified in neurons, in mediating motor function and neuronal connectivity [85, 86]. Whilst it has been previously postulated that therapeutic strategies based on PDI may be useful in ALS and possibly other protein misfolding disorders [87], the current study shows that PDI is also protective against DNA damage, another mechanism more recently associated with ALS pathophysiology. Thus, these findings broaden these earlier findings and reveal that PDI has wider applicability to ALS than previously recognised.

In conclusion, here we demonstrate a novel protective role for PDI in the nucleus against cellular DNA damage, induced by etoposide, H_2_O_2_ and mutant TDP-43. Therefore, novel therapeutic strategies involving PDI may have potential for ALS and other conditions involving the accumulation of DNA damage.

---

**Supplementary Figure 1.**
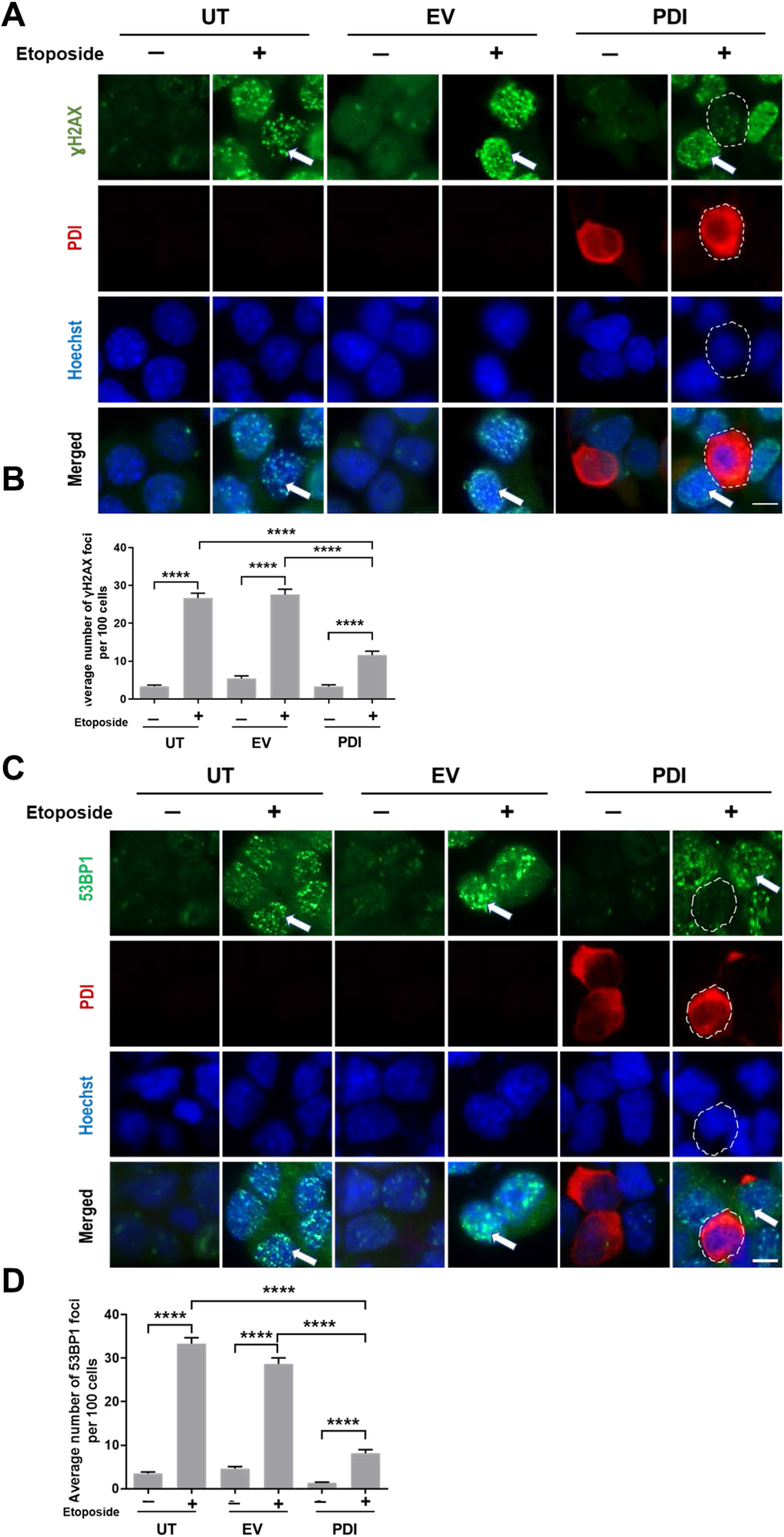
PDI inhibits the formation of DNA damage foci induced by etoposide in NSC-34 cells. **(A, C)** NSC-34 cells overexpressing PDI tagged with V5 (red) or empty vector pcDNA3.1(+) were subjected to immunocytochemistry following treatment with 13.5 μM etoposide for 30 minutes at 24 hours post-transfection, using anti-**γ**H2AX (A) or anti-53BP1(C)(both green) and V5 antibodies (red). Nuclei were stained with Hoechst (blue). Arrows represent **γ**H2AX foci(A) 53BP1 foci(C); Scale bar: 10 μm. **(B, D)** Quantification of the average number of foci per 100 cells shown in (A, D); PDI over-expression significantly reduced the formation of **γ**H2AX and 53BP1 foci in NSC-34 cells, following etoposide-induced DNA damage, ****p ≤0.0001, one-way ANOVA followed by Tukey’s multiple comparison post-hoc test. All values represent mean ± SEM.

